# Causality, Transfer Entropy and Allosteric Communication Landscapes in Proteins with Harmonic Interactions

**DOI:** 10.1101/084764

**Authors:** Aysima Hacisuleyman, Burak Erman

## Abstract

A fast and approximate method of generating allosteric communication landscapes is presented by using Schreiber's entropy transfer concept in combination with the Gaussian Network Model of proteins. Predictions of the model and the allosteric communication landscapes generated show that information transfer in proteins does not necessarily take place along a single path, but through an ensemble of pathways. The model emphasizes that knowledge of entropy only is not sufficient for determining allosteric communication and additional information based on time delayed correlations has to be introduced, which leads to the presence of causality in proteins. The model provides a simple tool for mapping entropy sink-source relations into pairs of residues. Residues that should be manipulated to control protein activity may be determined with this approach. This should be of great importance for allosteric drug design and for understanding the effects of mutations on protein function. The model is applied to determine allosteric communication in two proteins, Ubiquitin and Pyruvate Kinase. Predictions are in agreement with detailed molecular dynamics simulations and experimental evidence.

**Significance:** Proteins perform their function by an exchange of information within themselves and with their environments through correlated fluctuations of their atoms. Fluctuations of one atom may drive the fluctuations of another. Information transmitted in this way leads to allosteric communication which is described as the process in which action at one site of the protein is transmitted to another site at which the protein performs its activity. Disruption of allosteric communication by mutation for example leads to disease. The present paper incorporates information theoretic concepts into the well known Gaussian Network Model of proteins and allows for rapid characterization of allosteric communication landscapes for normal functioning as well as malfunctioning proteins.

## Introduction

Transfer of entropy from one subsystem of a protein to another is now becoming a subject of interest because of its relation to information flow and allosteric communication. Allosteric communication is the process in which action at one site of a protein is transmitted to another site at which the protein performs its activity. Protein-protein and protein-DNA interactions, drug action and all processes that depend on signal transduction involve allosteric activity for the system to carry out its normal function. Most known cancer causing mutations lead to the disruption of normal allosteric communication. Recent findings show that allosteric activity is entropic in nature and depends on information transfer from one part of the protein to the other [1, 2] through coordinated fluctuations of residues. Transmission of effects through correlated fluctuations is a universal property of all proteins and not only of allosteric ones. In this sense all proteins may be regarded as intrinsically allosteric in nature [3]. This new view freed the understanding of allostery from the limited picture of discrete two state transitions and opened a broader vista in terms of entropy transfer in proteins. The idea of transfer entropy, recently introduced by Schreiber [4], is the appropriate one for understanding information flow and communication in proteins. van der Vaart applied the Schreiber equation to determine information flow between Ets–1 transcription factor and its binding partner DNA [5], Barr et. al., [6] quantified entropy transfer among several residues in a molecular dynamics analysis of mutation effects on autophosphorylation of ERK2, Corrada et. al., [7] analyzed entropy transfer in antibody antigen interactions, Zhang et. al., [8] applied the method to understand changes in correlated motions of the Rho GTPase binding domain during dimerization. Presently, one of the requirements for calculating entropy transfer in proteins is to run molecular dynamics simulations in the order of a microsecond. Considering the urgent need for determining information transfer in malfunctioning proteins, the molecular dynamics technique becomes a serious bottleneck and a rapid characterization is needed. The aim of this paper is to provide a rapid scheme of computing entropy transfer in proteins and show that its results agree with detailed molecular dynamics based predictions. For this purpose we formulate transfer entropy using the Gaussian Network Model (GNM) which is based on harmonic interactions between contacting pairs of residues [9]. Calculation times for determining transfer entropy between all pairs of residues of proteins, even of extremely large protein complexes, using GNM can now be performed in the order of seconds on a laptop computer.

## Results of the Model

Using the GNM version of transfer entropy developed in this paper, we studied two important proteins, Ubiquitin and the Pyruvate Kinase tetramer.

Ubiquitin, (PDB code 1UBQ), is a 76 amino acid protein. It consists of 8 distinct secondary structures that actively take part in interactions with a large number of proteins. Ubiquitin, although not known as an allosteric protein itself, communicates with several other proteins to send information from one binding partner to another, and hence can be regarded as having intrinsic allosteric properties [10]. Recently, we used a 600 ns molecular dynamics trajectory to determine communication patterns in Ubiquitin resulting from entropy transfer [11]. Here, we compare the results of the present model with those from molecular dynamics (MD) simulations. The values of *T_i_*_→□_ (*τ*) for Ubiquitin from Eq.(9) are shown by the lower curve indicated as GNM in Fig 1. The upper curve shows the results of MD simulations. Both curves are obtained for the entropy associated with the alpha carbons of the residues. The GNM data is scaled and translated relative to the MD data for easier comparison. The secondary structures of Ubiquitin are shown in the upper part of the figure, where α and β stand for alpha helix and beta strand, respectively. The turns between the secondary structures are not indicated in the figure. The GNM and MD curves show good agreement in that the residues with high entropic activity are common to both except the β5 strand and the loop between β4 and β5.

**Fig 1.**
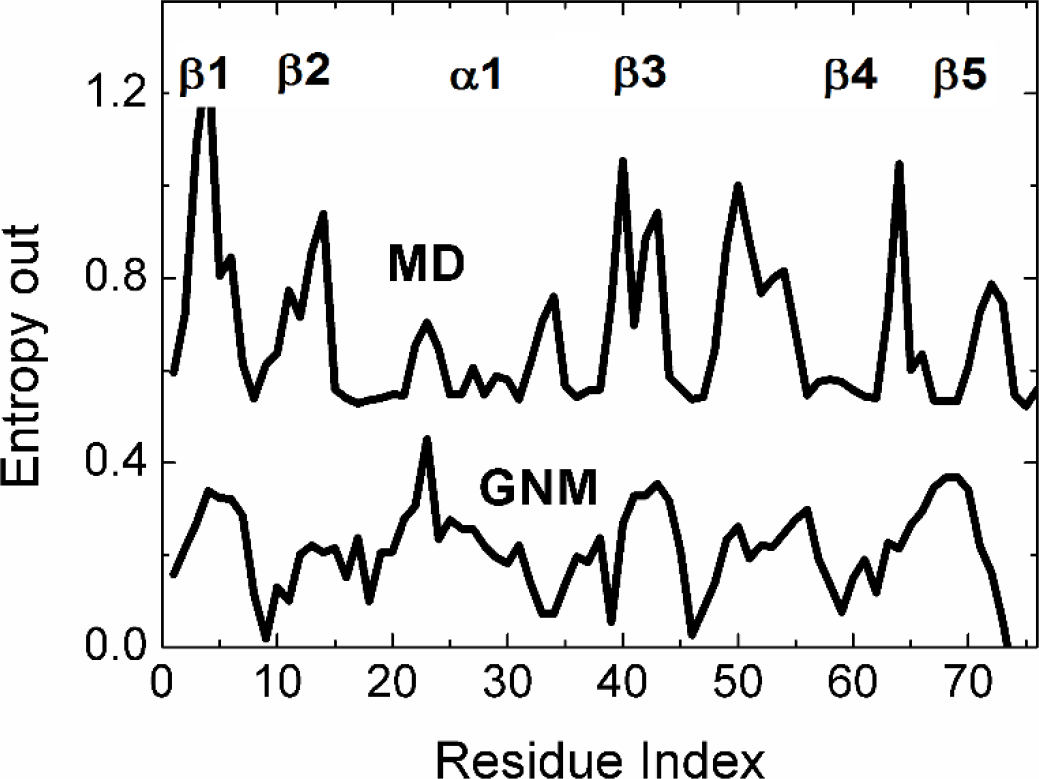
Entropy results of each alpha carbon of Ubiquitin. Upper curve is the entropy result calculated with MD data and lower curve is the entropy result calculated with GNM.

For a more detailed analysis we present, in Fig 2, the transfer entropy values *T_i_* _→ *j*_ (*τ*) for each pair of residues of the protein.

**Fig 2.**
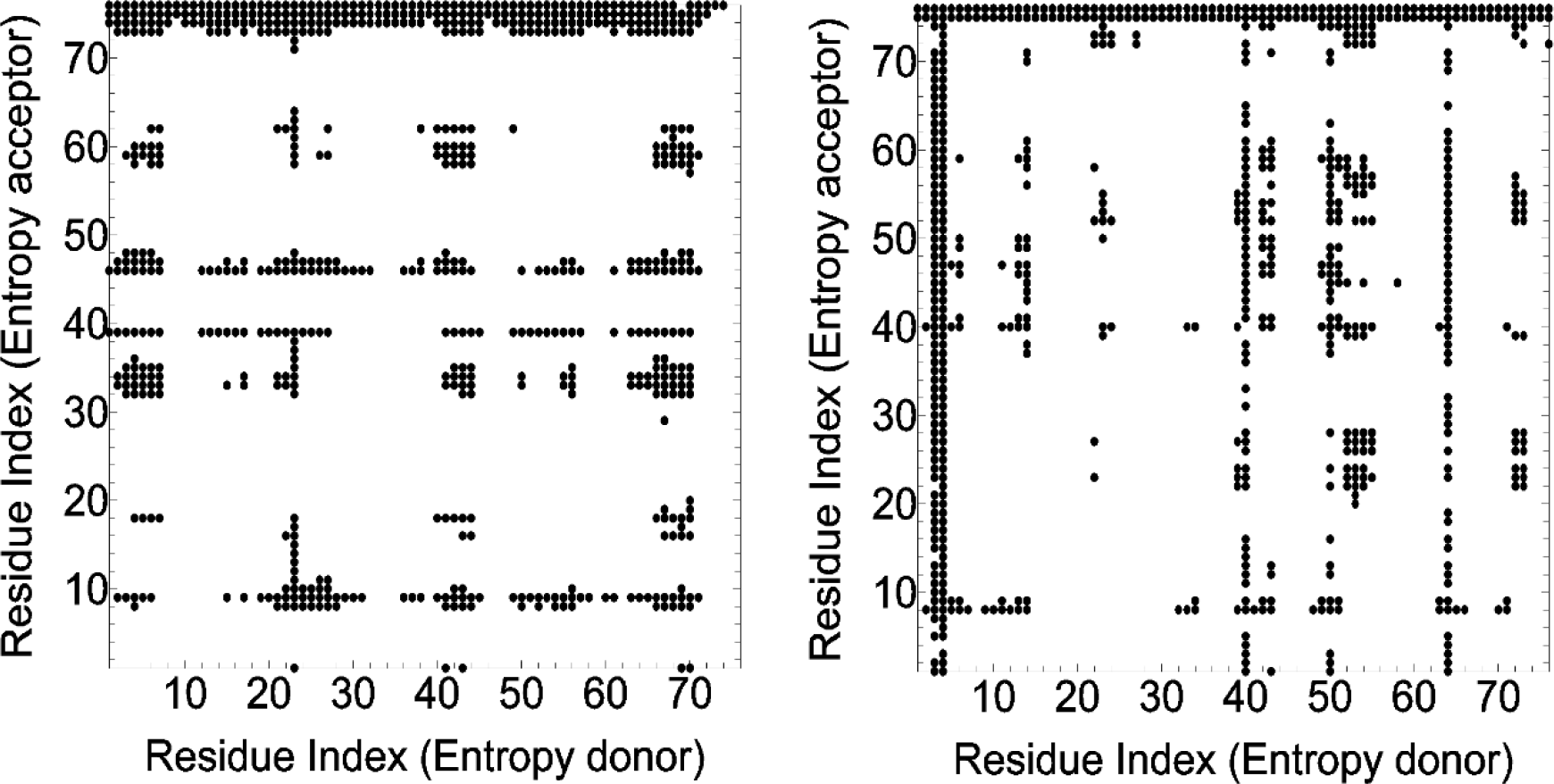
Transfer entropy between Ubiquitin residue pairs. Left panel shows GNM results and right panel shows MD results. Entropy is being transferred from the donor(x axis) to the acceptor(y axis).

In this figure, we compare the entropy transfer values *T_i_* _→ *j*_ (*τ*) calculated from Eq. (8) of Ubiquitin obtained by GNM (left panel) and from MD results of Reference [11] (right panel). The value of *τ* is taken such that relaxations decay to 1/e of the initial [A1]values. The horizontal axes denote the indices of residues that act as entropy donors. The vertical axes denote indices of residues that act as residue acceptors. For example the horizontal line of points corresponding to residue 76 in the ordinate of both panels indicates that this residue is an entropy sink and takes the entropy of all other residues. Several such entropy acceptors are common to both panels in the figure. In this respect, residues L8, Q40, G47, N60 and G75 absorb entropy from several residues of UBQ. An example to an entropy donor residue is I23. On the left panel, a sequence of points perpendicular to the x axis at 23 indicates that I23 gives entropy to several other residues. There is a similar trend for residue I23 shown by the MD data in the right panel. The vertical direction at 23 here is not as densely populated as the corresponding one on the GNM data, nevertheless there is a large set of residues that accept entropy from I23. According to MD results, residue I3 is an entropy donor to almost all other residues, as indicated by the vertical points at 3. GNM results show that residues I3-V5 are entropy donors to several residues but not exactly as shown by the discrete succession of points in the MD results. Residue Q40 shows also approximately similar behavior as entropy acceptor for both GNM and MD. Other residues with common features for GNM and MD data may also be seen from the two panels of Fig 2.

In Fig 3, we present the surface plot of the antisymmetric part of Eq. (10) for Ubiquitin. In this figure, the red regions show entropy transfer from the entropy donor to the entropy acceptor residue, identified on the abscissa and the ordinate, respectively. The blue regions show entropy transfer to the donor residue. If a pair of residues, i from the entropy donor axis and j from the entropy acceptor axis leads to a positive value on the surface, then i provides entropy to j and vice versa for the negative surface. Thus, the negative blue trough for i=76 on the entropy donor axis means i=76 extracts entropy from all residues along the trough. The surface depicted in Fig 3 gives a view of the communication landscape of UBQ.

**Fig 3.**
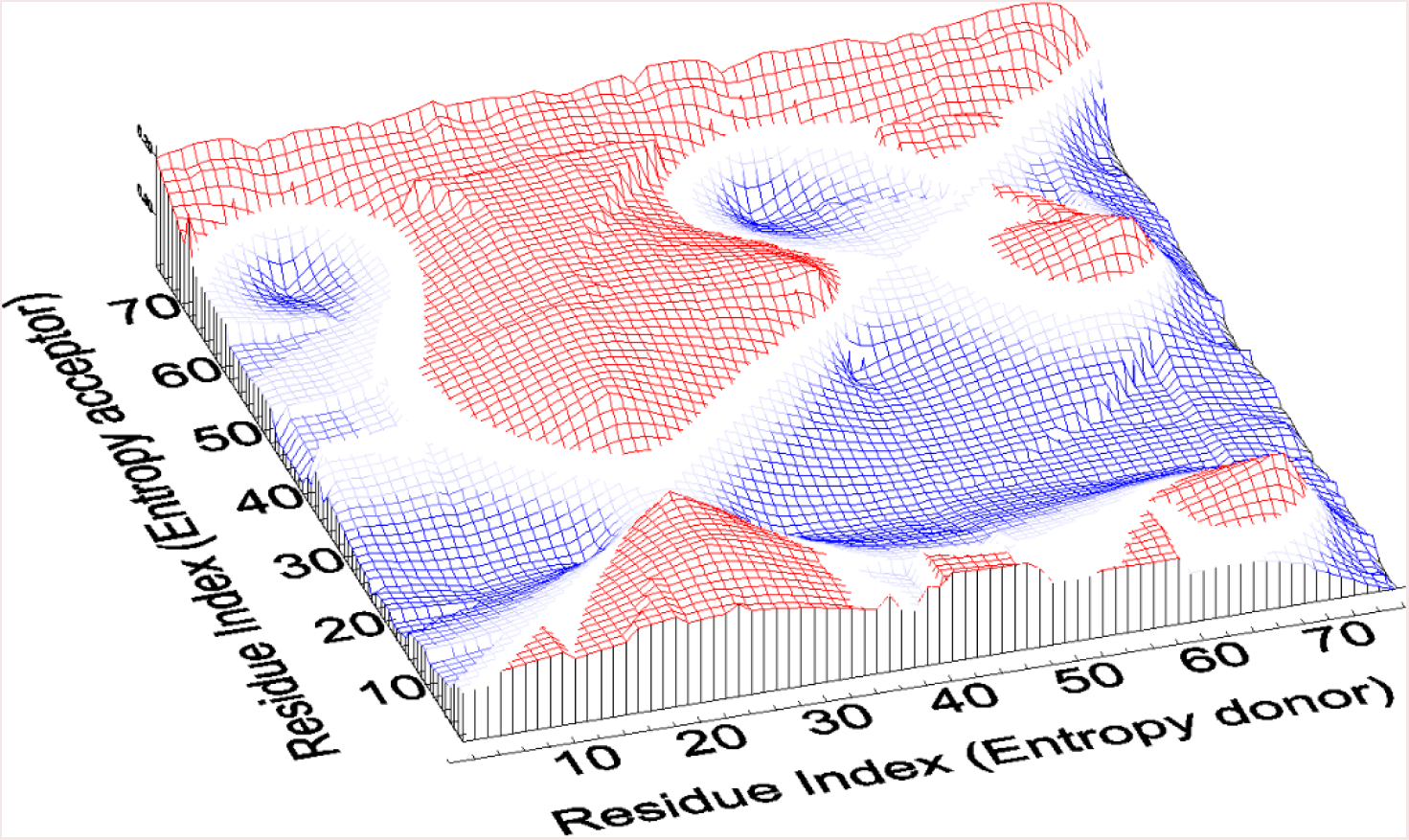
Surface plot of entropy transfer in Ubiquitin. Red regions denote entropy transfer from the entropy donor axis to entropy acceptor axis. Blue regions show the reverse, entropy transfer into the residue identified on the entropy donor axis.

A more concise view of sink and reservoir residues may be seen from the net entropy out from a residue i curve obtained by Eq. (11). In Fig 4 results for UBQ are shown.

**Fig 4.**
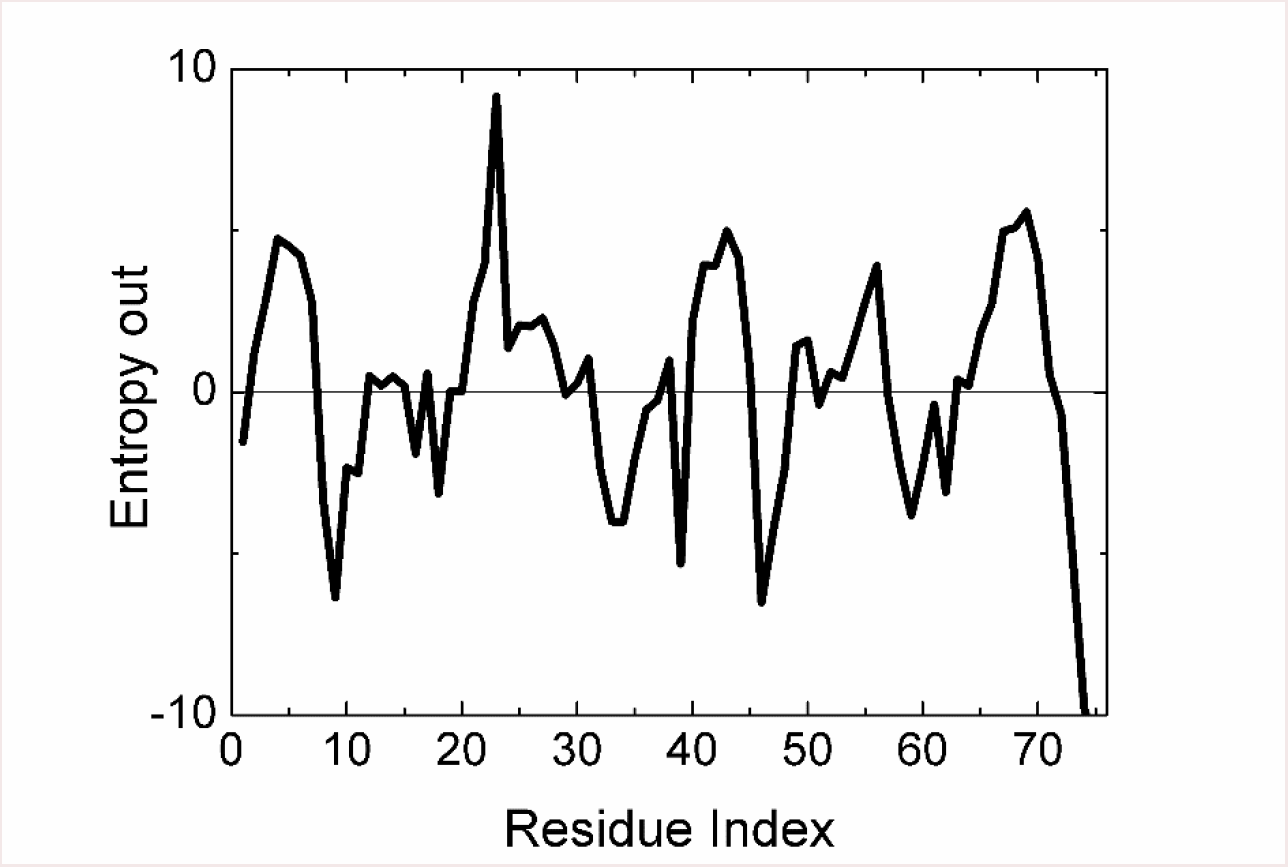
Net Entropy out from residues of UBQ.

The negative peaks indicate that the corresponding residue acts as an entropy sink, extracting entropy from the protein. The positive peaks are the entropy reservoirs that provide entropy to the protein. An interesting feature from the figure is that the two C-terminal residues G75 and G76 act as residue sinks, absorbing the entropy of the remaining residues of the protein.

## Pyruvate Kinase

Pyruvate Kinase from *Leishmania mexicana* (PDB code 3HQQ), which is a homotetrameric enzyme having 1992 amino acids in total and 498 amino acids per protomer, each consisting of 28 secondary structures and 4 domains named as N-terminal(residues between 1-17), A(residues between 18-88 and 187-356), B(residues between 89-186) and C(residues between 357-498). It catalyzes the last step in glycolysis and is known to be allosterically activated by the binding of fructose biphosphate (FBP). Ligand free and allosterically activated states of PK differ in rigid body rotations of the AC core. This rotation results in R310 making hydrogen bonds with R262 and G263 on an alpha helix and this helix unwinds in the absence of FBP. R310 plays a crucial role because it shows the greatest conformational changes between the inactive and active states of the enzyme, but it has been proven that it is necessary but not sufficient for allosteric transition. FBP interacts with E451 and G487 in the C–domain and affects the motion of the B-domain by increasing the rigidity of the enzyme and consequently its activity although it binds to a site over 40 Å away from the active site [12-14]. Here we used the alpha carbons of the crystal structure of apo form of PK to determine communication patterns between its residues with GNM based calculations by using Eq (8).

Entropy transfer and allosteric activity in PK takes place at two different scales. Firstly, transfer among the four protomers, and secondly within each protomer. We analyze them separately. In Fig 5, we present the communication landscape for the tetramer. Red regions denote positive and blue regions denote negative entropy transfer from the entropy donor to acceptor residue pairs. We see a continuous negative trough for R310 of the entropy donor axis. A negative entropy donor means R310 extracts entropy from other residues of the protein including all other protomers. R310 makes contact with G263 and G266 of another protomer through its large interface. This leads to a long range entropy transfer inclusive of all four protomers. The red peaks between S1 and A20 of the entropy donor axis indicates that these residues provide entropy to the rest of the protein. These peaks extend over all other protomers and hence represent long range entropy transfer. Inspection of the crystal structure shows that these residues belong to the short helix hanging on a long loop near the short interface. Thus each protomer provides entropy to other protomers through a certain set of residues and extracts entropy from other protomers through another set of residues. This constitutes the basis of the mechanism of the inhomogeneous rocking motions of the tetramer. The communication landscape of Fig 5 is essentially a pictorial representation of the large scale rocking motions. The symmetry of rock and lock motion across all four protomers is noticeable from the figure.

**Fig 5.**
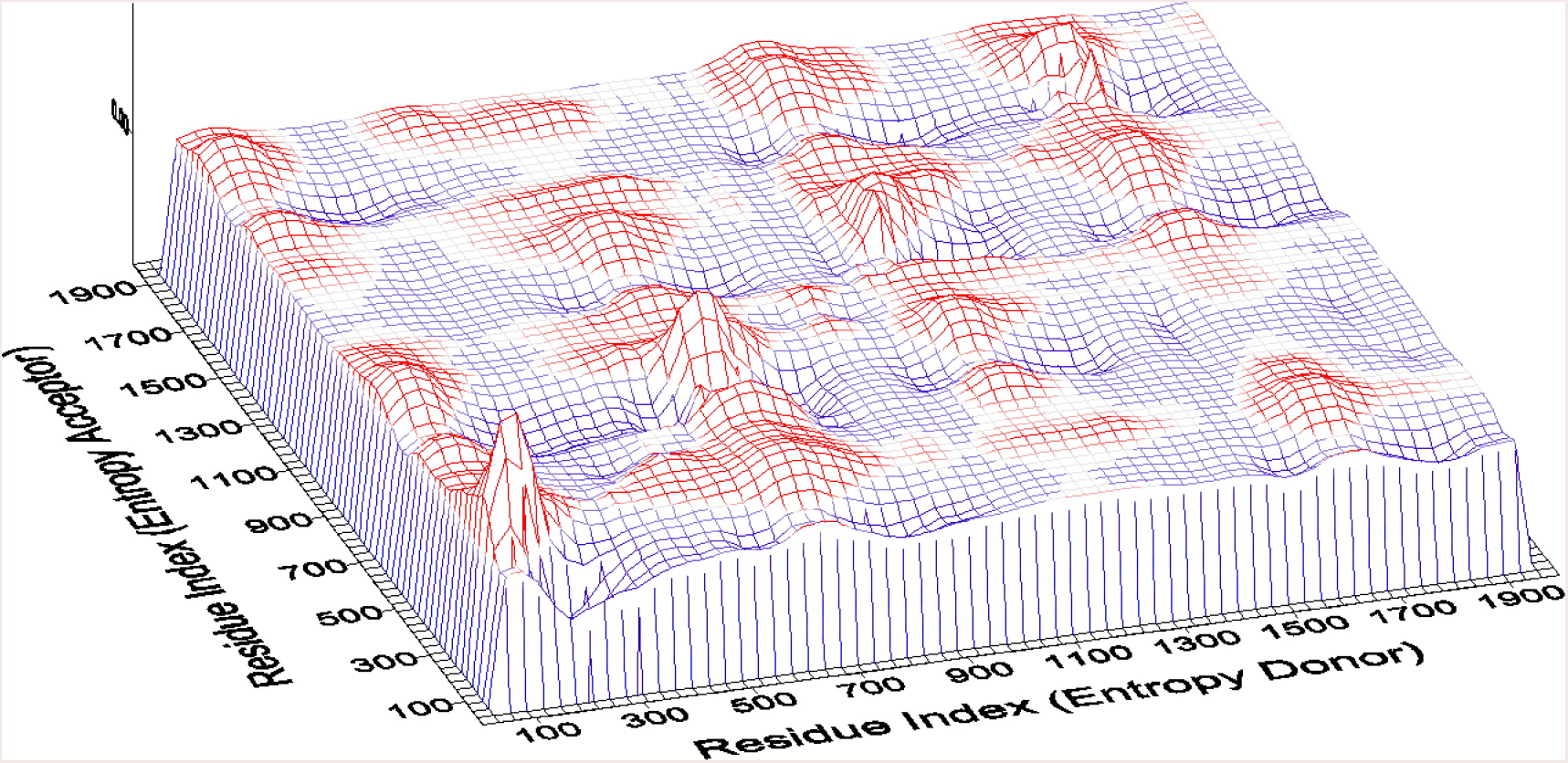
Transfer entropy between Pyruvate Kinase residue pairs. Red regions denote positive and blue regions denote negative entropy transfer between residue pairs.

In the interest of understanding intra-residue communication in a more detailed way i.e., transfer within a protomer, we present entropy transfer over short scales in Fig 6. The regions corresponding to entropy donor residue indices between 350 and 498 are predominantly blue, showing that these residues extract entropy from the remaining residues. The red domains corresponding to the entropy donor residues A70 to T170 are the residues that give their entropy to the others. FBP binds to residues L399, S400, N401, T402, S405, H480, A481 and G487, which are all in the blue region of Fig 6, extracting entropy from the red regions, i.e., residues between A70 and T170.

**Fig 6.**
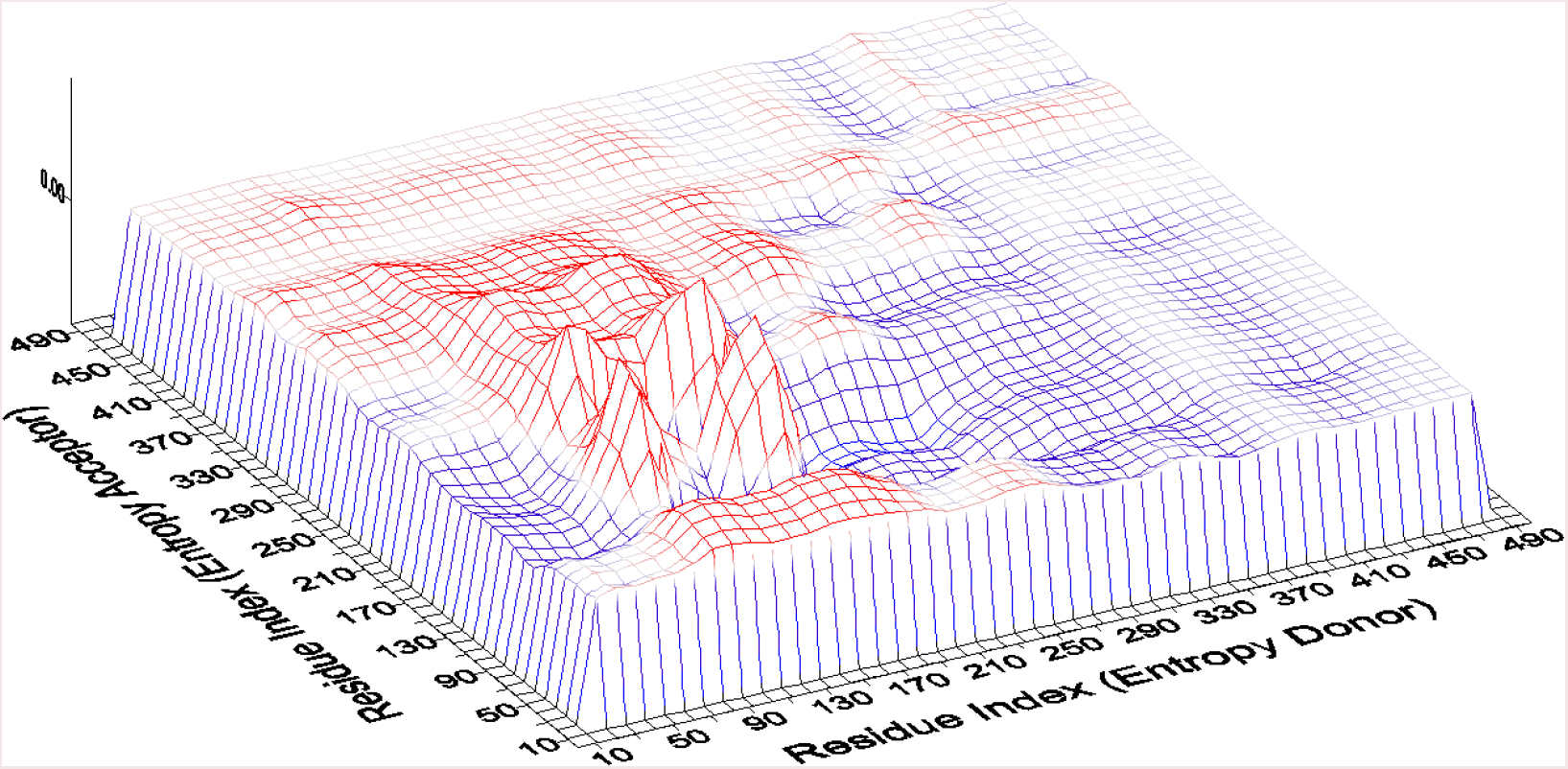
Transfer entropy between residue pairs of one PK protomer. Red regions denote positive and blue regions denote negative entropy transfer between residue pairs.

Fig 7 shows the three dimensional structure of a protomer. The green molecule is FBP. Highlighted blue residues around FBP are the binding residues that fall in the blue region of Fig 6. They extract entropy. The blue ribbon and the highlighted blue head in the figure are the residues A70-T170 that donate entropy to others.

**Fig 7.**
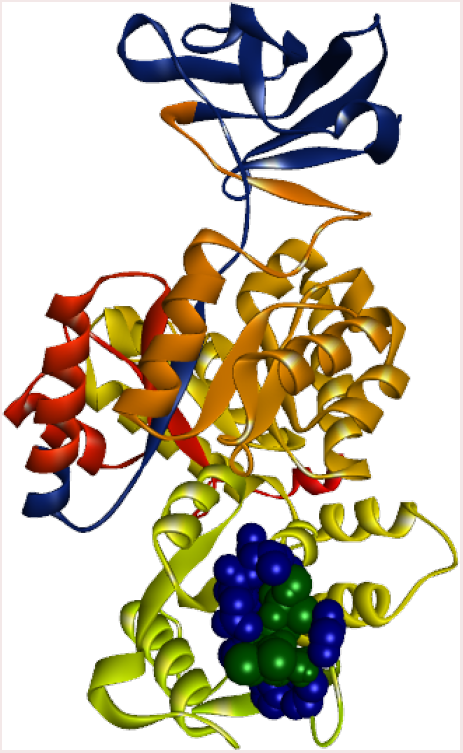
Cartoon of a single protomer of PK in complex with FBP. Blue residues in ribbon representation (A70-T170) donate entropy to others, green molecule is FBP and blue residues in CPK representation are the binding residues of FBP.

## Discussion

In this paper we present a fast but approximate method of determining allosteric communication landscapes in proteins using the Gaussian Network Model which is based on harmonic interactions between contacting residues. The method is based on the transfer entropy concept introduced recently by Schreiber, and shows that knowing only the energy landscape is not sufficient to predict information transfer and allosteric communication, and that time delayed correlations are necessary. The allosteric communication landscapes presented in Figs. 3, 5 and 6 show that information transfer in proteins does not necessarily take place along a single path, but over an ensemble of pathways. The model also emphasizes that knowledge of entropy only is not sufficient for determining allosteric communication and additional information based on time delayed correlations has to be introduced, which leads to the presence of causality in proteins. The possibility of causality in proteins allows for identifying driver-driven relations for pairs of residues. The GNM method of entropy transfer provides a rapid tool for determining the allosteric communication landscape for proteins. Construction of a landscape for a protein as large as 2000 residues now takes less than one minute on a desktop. We performed a comparative analysis of allosteric communication between residue pairs in Ubiquitin and Pyruvate Kinase with the present method and showed that the results are in good agreement with molecular dynamics based predictions. Evaluating the communication map for the Pyruvate Kinase by molecular dynamics would take several months on a supercomputer. With the GNM approach, we were able to determine the allosteric communication landscape of the 1992[A2] residue protein Pyruvate Kinase. The results agreed with experimentally known allosteric communication features of the complex. The GNM entropy transfer model provides a simple tool which maps the entropy sink-source relations into pairs of residues. By this approach residues that should be manipulated to control protein activity may be determined. This should be of great importance for allosteric drug design and for understanding effects of mutations on protein function.

## Methods

Following Schreiber's work [4], we write the transfer entropy *T_i_* _→ *j*_ (*τ*) from the fluctuation trajectory Δ*R_i_* (*t*) of atom i to that of j Δ*R* _*j*_ (*t* + *τ*) at time *t* +*τ* as

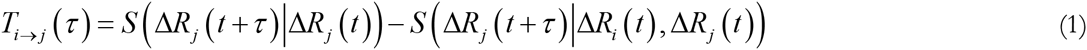

The conditional entropies on the right hand side of the Schreiber equation for two events Δ*R_i_* (*t*) and Δ*R* _*j*_ (*t* +*τ*) in a steady state process is given in Boltzmann units, i.e., the Boltzmann constant taken as unity by

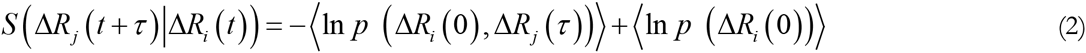

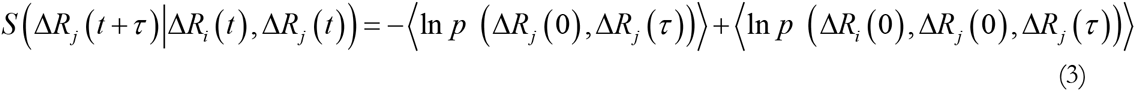

Detailed derivations of the expressions presented in this paper are provided in the Supplementary Material. In the coarse graining approximation we focus only on the alpha carbon of each residue.

Substituting Eqs. (2) and (3) into Eq. (1), leads to

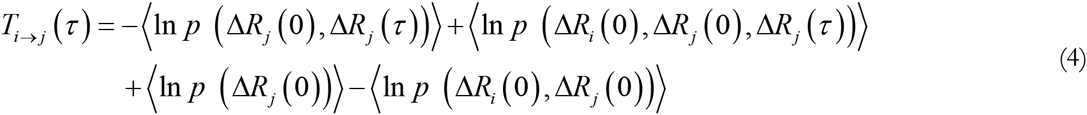

The Gaussian Network Model is characterized by the spring constants matrix Γ where a spring of constant unity is assumed between residues in contact. It is defined as follows: Γ_*ij*_ equates to–1 if alpha carbons of residues i and j are within a cutoff distance of *r_C_* and to zero otherwise. Each ith diagonal element Γ_*ii*_ is equal to the negative sum of the ith row. The time correlation of fluctuations is given by the GNM as [15]

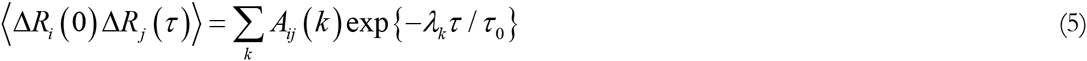

Where

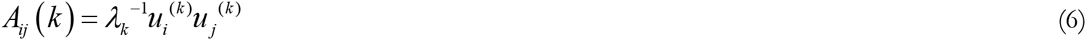

with *λ_k_* being the kth eigenvalue and *u_i_*^(*k*)^ being the ith component of the kth eigenvector of the Γ matrix. The probability distribution of a Gaussian of n variables, Δ*R* = [ Δ*R*_1_, Δ*R*_2_, Δ*R*_3_, &, *n*] is

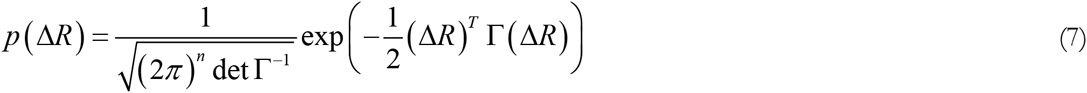

where, Γ^−1^ is the matrix of covariances 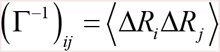

Taking the logarithm of Eq. (7) and averaging and substituting into Eq. (4) leads to the final expression for entropy transfer in the GNM as:

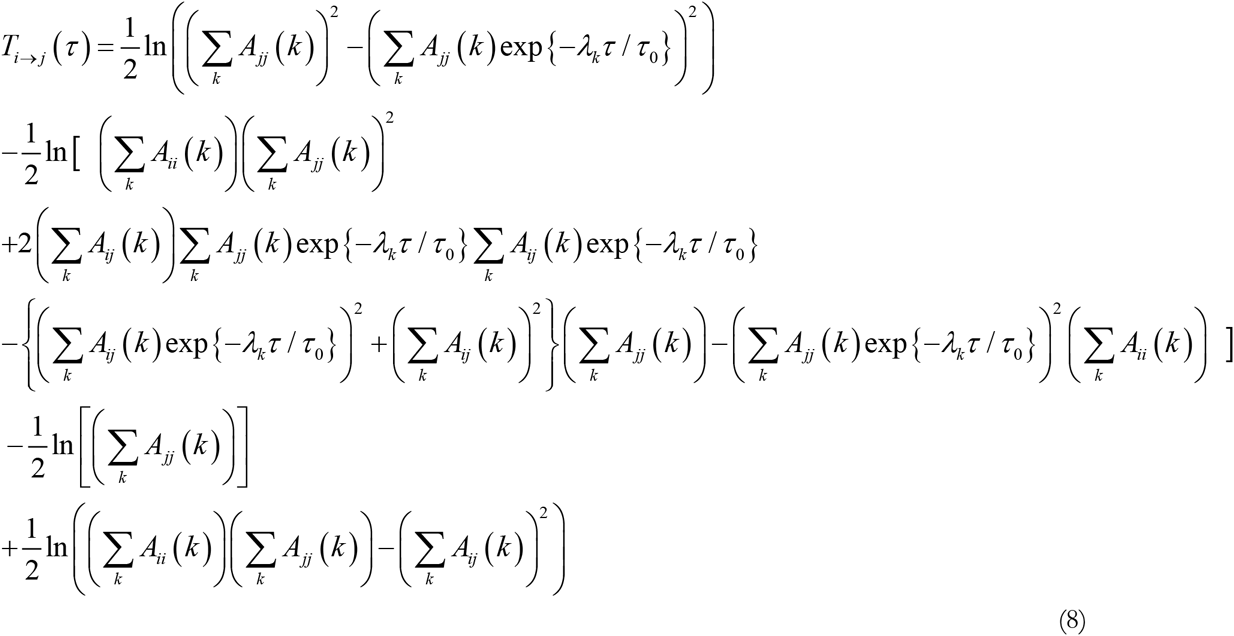

The net entropy transfer *T_i_*_→ □_ (*τ*) from residue i to the rest of the protein is obtained by summing

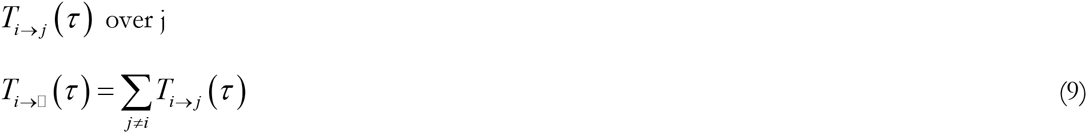

Defined in this way, *T_i_*_→□_(*τ*) is a measure of entropic activity of residue.j[A3]

The matrix *T_i_* _→ *j*_ (*τ*)may be written as the sum of its symmetric and antisymmetric parts as

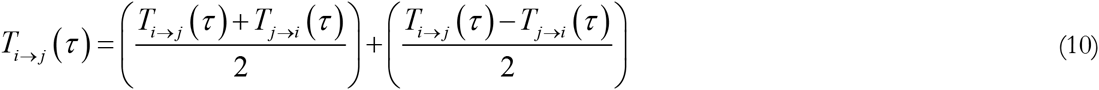

The second parenthesis on the right hand side is nonzero due to causality of events. If the fluctuations of residue i drive the fluctuations of residue j stronger than the action of j on i, then residue i acts as the driver and j as the driven. The antisymmetric part has both negative and positive components, corresponding to entropy sinks and entropy reservoirs. Summing up the right hand side over j gives the net entropy going out of residue i as:

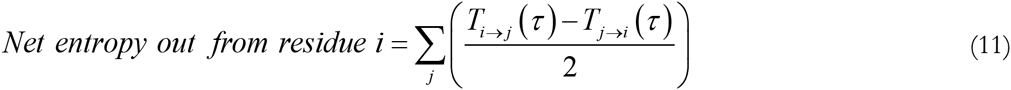

The time delay *τ* that appears in the equations depends on the spring constant of the harmonic interactions. Molecular dynamics simulations of proteins at physiological temperatures shows that fluctuations of two residues i and j are in the order of 5 nanoseconds [11, 16]. However, it is not possible to establish an exact quantitative correspondence between the GNM values and real time parameters. In all the calculations below, we took *τ* as the time for which correlations decay to 1/e of their initial values.

## Supplementary material for the derivation of equations in the main manuscript

We assume that the trajectories, Δ*R_i_* (*t*) and Δ*R* _*j*_ (*t*) for two atoms are known. We now consider two events separated in time by *τ*, with the condition that Δ*R_i_* coming before Δ*R_j_*. The conditional entropy for these two events is defined by

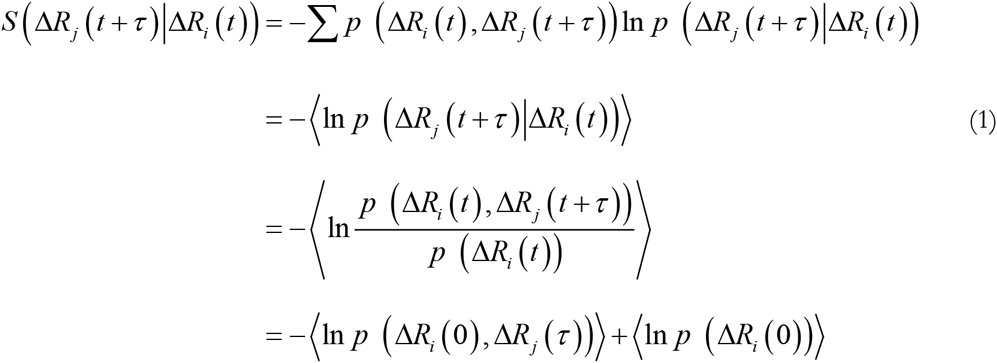

where, the summation is over all states for i and j, and the condition of stationarity is used in the last equation. Throughout the paper we take the Boltzmann constant as unity.

Following Schreiber's work [1], we write the transfer entropy *T_i_* _→ *j*_ (*τ*) from trajectory i to j at time τ as

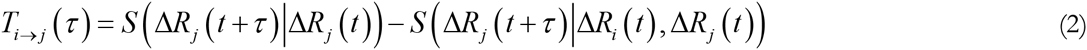

Using the relations for conditional entropy, this may be written as

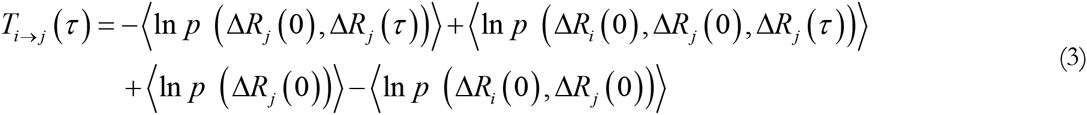

The probability distribution of a Gaussian of n variables, Δ*R* = [ Δ*R*_1_, Δ*R*_2_, Δ*R*_3_, &, *n*] is

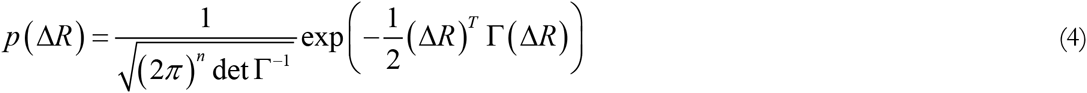

where, Γ_*ij*_ is defined from the Gaussian Network Model (GNM) [2] as the matrix of spring constants which has the value–k if atom i is within a given cutoff distance of *r_C_* which is usually taken as 7 Ångstroms. If the distance between two atoms is larger than *r_C_*, then Γ_*ij*_ equals zero. The diagonal elements Γ_*ii*_ is equal to the negative sum of the ith row. The magnitude of the spring constant k is immaterial, and hence is taken as unity. Γ^−1^ is the matrix of covariances with 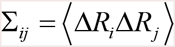.

Then,

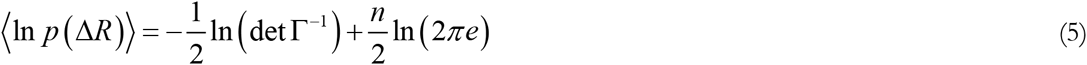

Each term in Eq. 3 now takes the following form

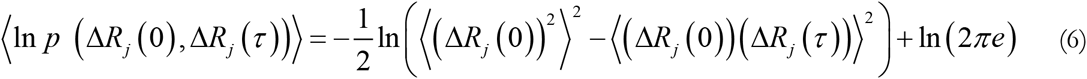

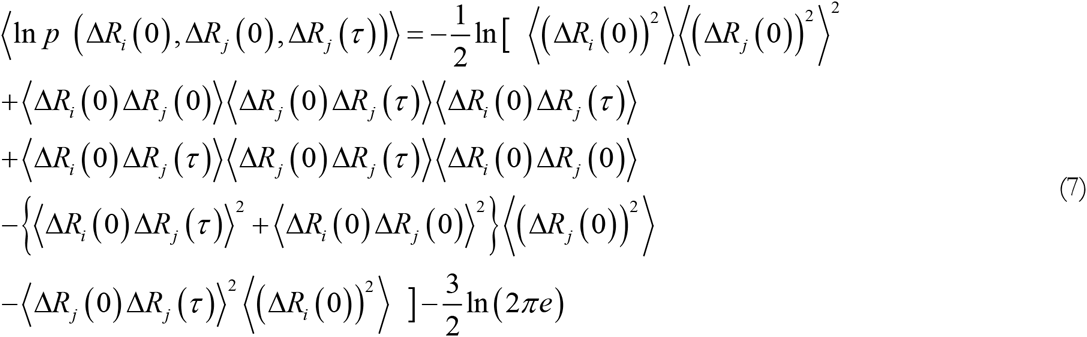

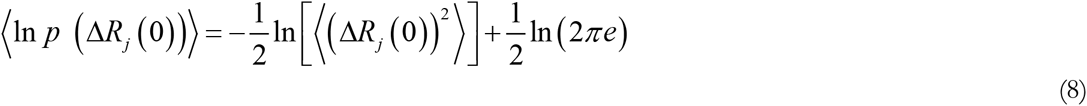

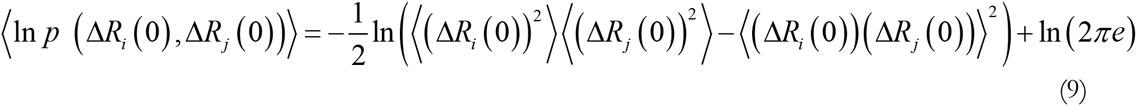

The time correlation of fluctuations according to the GNM is given as [3]

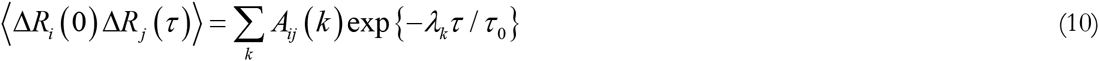

where

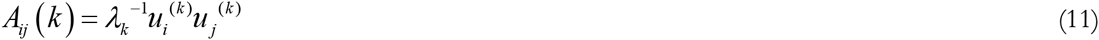

with *λ_k_* being the kth eigenvalue and *u_i_*^(*k*)^ being the ith component of the kth eigenvector. For *τ* = 0,

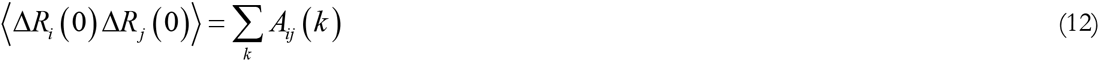

Substituting these into into Eqs. 6-9 and then into Eq 3 leads to

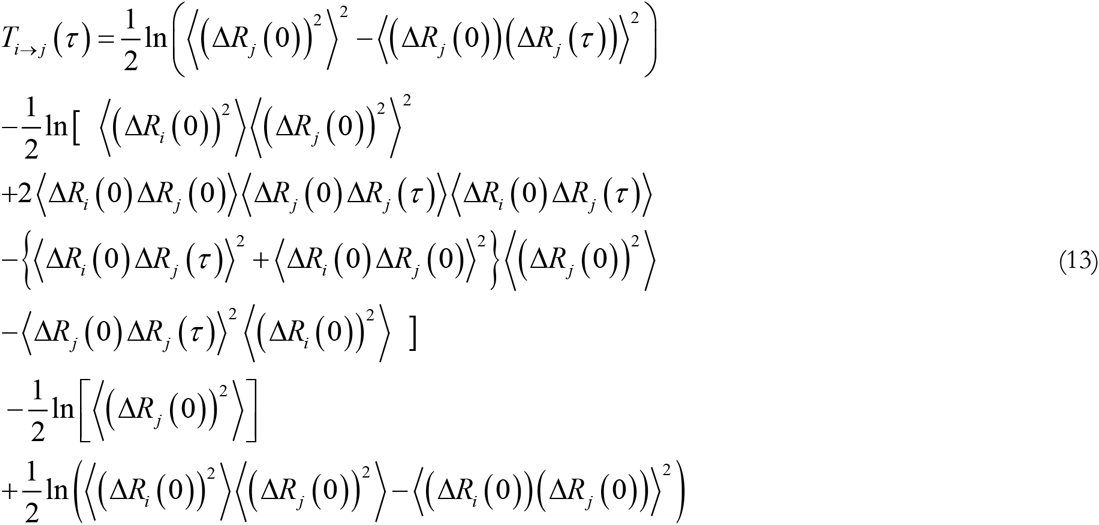

When the fluctuations of the two residues are uncorrelated, this quantity becomes zero. Substituting the eigenvalues and eigenvectors into Eq. 13 leads to the final expression for entropy transfer

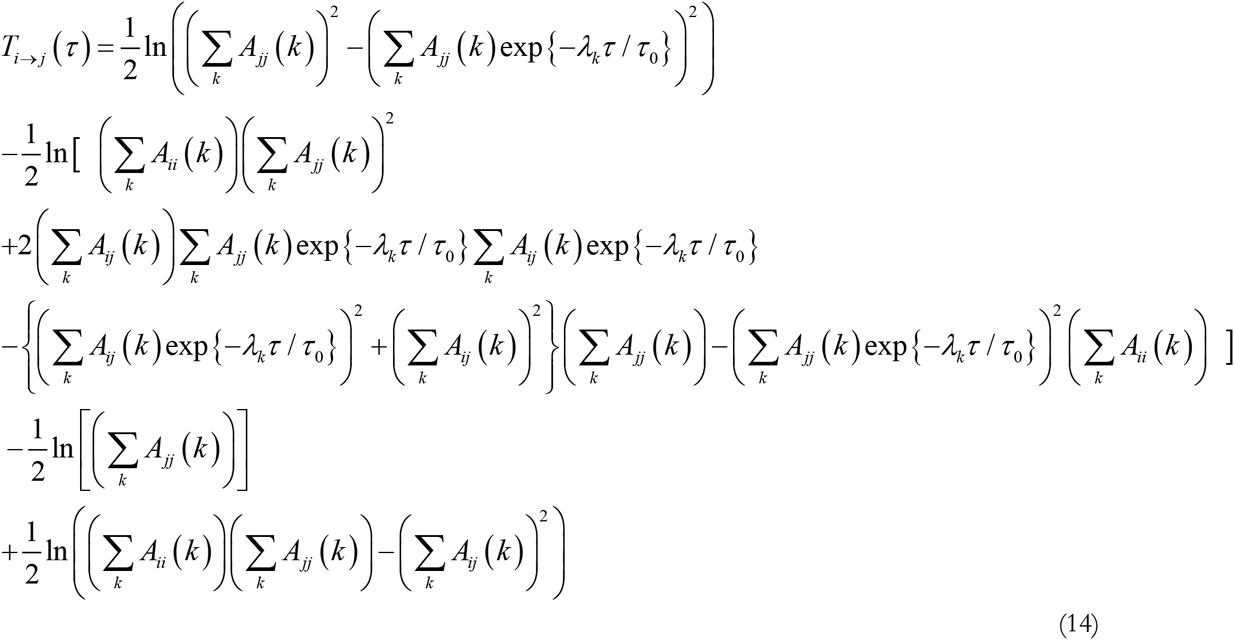

